# A high-throughput metabolic microarray assay reveals antibacterial effects of black and red raspberries and blackberries against *Helicobacter pylori* infection

**DOI:** 10.1101/2020.12.14.422663

**Authors:** Candace Goodman, Katrina Lyon, Aitana Scotto, Mandi M. Roe, Farimah Moghimpour, T. Andrew Sebrell, Andrew B. Gentry, Ganesh Bala, Gary Stoner, Diane Bimczok

## Abstract

*Helicobacter pylori* is an important bacterial pathogen that causes chronic infection of the human stomach, leading to gastritis, peptic ulcer disease and gastric cancer. Treatment with appropriate antibiotics can eliminate *H. pylori* infection and reduce the risk for severe disease outcomes. However, since *H. pylori* is becoming increasingly resistant to standard antibiotic regimens, novel treatment strategies are needed. Previous studies have demonstrated that black and red berries may have antibacterial properties. Therefore, we analyzed organic extracts and powders from black and red raspberries and blackberries and determined their antibacterial effects on multiple *H. pylori* strains. We used high-performance liquid chromatography to measure berry anthocyanins, which are considered the major active ingredients. To monitor antibiotic effects of the berry preparations on *H. pylori*, we developed a high-throughput metabolic growth assay based on the OmniLog™ system. All berry preparations tested had significant bactericidal effects *in vitro*, with MIC_90_ values ranging from 0.49 to 4.17%. We next used human gastric epithelial organoids to evaluate biocompatibility of the berry preparations and showed that black raspberry extract, which had the strongest antimicrobial activity, was non-toxic at the concentration required for complete bacterial growth inhibition. To determine whether dietary black raspberry application could eliminate *H. pylori* infection *in vivo*, mice were infected with *H. pylori* and then were placed on a diet containing 10% black raspberry powder. However, this treatment did not significantly impact bacterial infection rates or gastric pathology. In summary, our data indicate that black and red raspberry and blackberry products have potential applications in the treatment and prevention of *H. pylori* infection, because of their antibacterial effects and good biocompatibility. However, delivery and formulation of berry compounds needs to be optimized to achieve significant antibacterial effects *in vivo*.

## Introduction

*Helicobacter pylori* is the major cause of human gastric disease worldwide[1, 2]. *H. pylori* is an acid-resistant, gram-negative bacterium that persistently infects the gastric mucosa of approximately half the world’s population, leading to chronic active gastritis [1]. A proportion of infected individuals also develop peptic ulcer disease, autoimmune gastritis or gastric adenocarcinoma, the second leading cause of cancer-related mortality [3]. In spite of decades of active research, no effective vaccine to prevent *H. pylori*-associated illnesses has been developed [4]. Once diagnosed, *H. pylori* infection is generally treated with a combination of antibiotics and proton pump inhibitors. However, increased resistance to two of the standard antibiotics included in *H. pylori* treatment regimens, clarithromycin and metronidazole, has been reported in multiple studies, with resistance rates ranging from 22-80% [5, 6]. Recently, clarithromycin-resistant *H. pylori* were included in the WHO’s high priority pathogens list for research and development of new antibiotics [7]. Moreover, poor patient compliance with complex medication regimens contributes to decreased treatment success [8, 9]. Therefore, eradication rates of *H. pylori* have dropped below 75% in several countries [10, 11]. The high failure rate of traditional *H. pylori* therapies points to an urgent need for novel alternative treatments or preventative strategies to combat *H. pylori* infection [12].

A significant body of research in recent years has shown that natural dietary components, especially plants, contain many bioactive compounds – neutraceuticals – with antibacterial effects [13–15]. Multiple different berries and their products show significant antimicrobial activity *in vitro* and *in vivo*, and some promising studies suggesting effectiveness against *H. pylori* have been published. Thus, data by Chatterjee et al. [16] showed significant inhibition of *H. pylori* growth in the presence of extracts from raspberry, strawberry, cranberry, elderberry, blueberry and bilberry. In another recent study, extracts from unripe Korean raspberries and elm tree bark used in combination significantly suppressed *H. pylori* growth both *in vitro* and in a mouse model [17]. In an *in vitro* model of *H. pylori* infection, cyanidin 3-O-glucoside, a key anthocyanin present in red and black berries, significantly decreased *H. pylori*-induced cell death [18]. Since berry products also have proven anti-inflammatory and anti-cancer effects, their potential application in *H. pylori* infection is particularly attractive.

In our study, we developed a high-throughput metabolic assay to screen different black raspberry, red raspberry and blackberry preparations for their ability to prevent *H. pylori* growth *in vitro.* We also used a gastric organoid model to evaluate biocompatibility and a mouse model of *H. pylori* infection to evaluate antimicrobial effects of black raspberry *in vivo*. Our results demonstrate that all berry powders and extracts tested caused significant reduction of *H. pylori* growth in two different strains at concentrations between 0.5 and 3%. An optimum preparation of black raspberry extract used at 0.5% led to complete inhibition of *H. pylori* growth but did not affect the viability of primary gastric epithelial cells. However, dietary application of black raspberry powder failed to achieve a significant reduction in *H. pylori* colonization in C57BL/6 mice infected with strain PMSS1. These results suggest that preparations from black and red raspberries and blackberries have potential as novel antimicrobial agents to combat *H. pylori* infection, but that preparation and application of the berry products need to be optimized to achieve antimicrobial activity *in vivo.*

## Materials and Methods

### Berry powders and preparation of extracts

Commercially available black raspberry (*Rubus occidentalis;* BRB), blackberry (*Rubus fruticosus;* BB) and red raspberry (*Rubus idaeus;* RRB) samples were either purchased as freeze-dried powders or as fresh-frozen whole berries. Fresh-frozen whole berries were processed into freeze-dried powder using a food processor and lyophilizer (**Figure 1a** and **Table 1**). Berry powders were either used directly in experiments (whole berries, WB, or powder) or processed for hexane/ethanol extraction (extracts, E). For hexane/ethanol extraction, nonpolar compounds were extracted using 3 × 100 mL hexane per 10 g powder. Suspensions were filtered between each extraction, and the hexane filtrate discarded. Anthocyanins and all water-soluble compounds were then extracted using an 80:20 ethanol:water mixture (3 × 100 mL per 10g sample). This extract was dried to a syrup under reduced pressure at 30°C then lyophilized (referred to as ethanol extract, E) to yield between 1-4 g per 20 g powder. All berry preparations were stored in airtight containers at −20° C until use.

**Figure 1.**
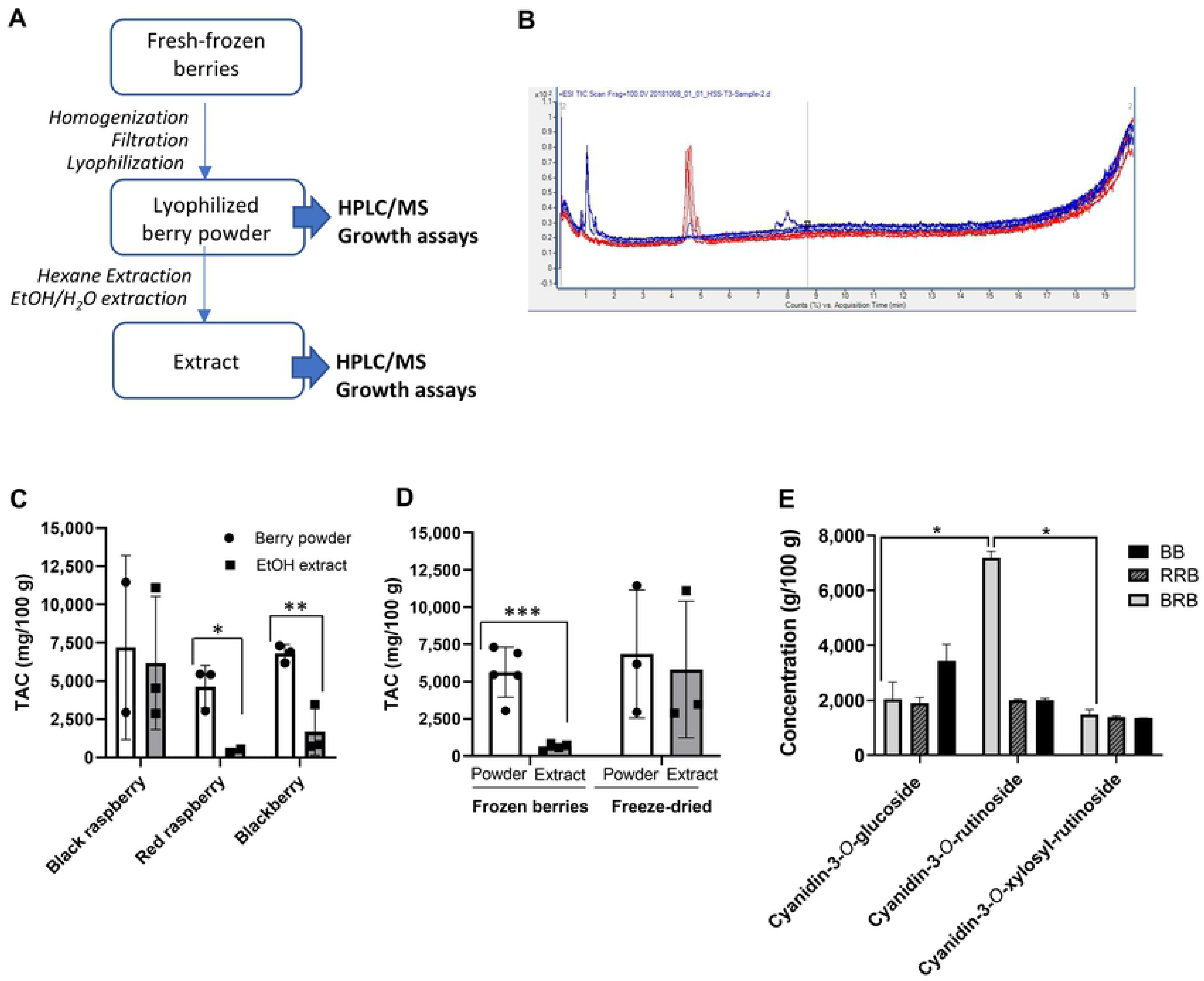
Preparation and anthocyanin content analysis of black raspberries, red raspberries and blackberries. (**A**) Workflow for berry preparation and analysis, (**B**) Representative LC/MS spectrum of berry preparations. Major peaks represent cyanidin-3-*O*-glucoside, cyanidin-3-*O*-rutinoside, and a combination of cyanidin-3-*O*-xylosylrutinoside and cyanidin-3-*O*-sambubioside. (**C**) Total anthocyanin content (TAC) in suspended powders and ethanol extracts of black raspberries (BRB), red raspberries (RRB) and blackberries (BB) determined by LC/MS. Individual data points, mean and SD are shown. (**D**) TAC in powders and extracts of BRB, RRB and BB purchased as fresh-frozen berries or as freeze-dried berry powder. Pooled data from all berries; individual data points, mean ± SD are shown. (**E**) Concentrations of major anthocyanins in suspended powders of BRB, RRB and BB. Statistically significant differences as determined by (**C, D**) Student’s t test (**E**) or 2-way ANOVA are shown as \**P*<0.05, \*\**P*<0.01 and \*\*\**P*<0.001.

**Table 1:**
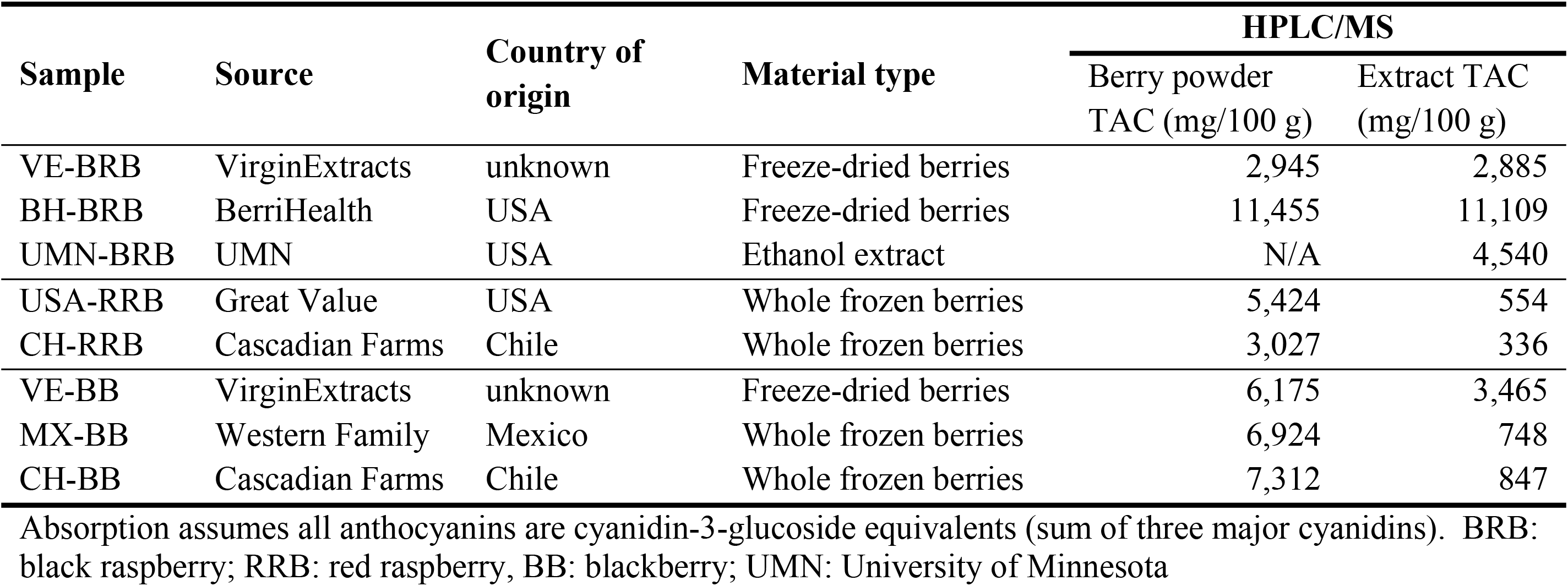
Total concentrations of anthocyanins in black and red raspberry and blackberry extracts determined by LC/MS.

### Analysis of anthocyanin content

Concentrations of major active compounds, i.e., cyanidin-3-*O*-glucoside, cyanidin-3-*O*-rutinoside, cyanidin-3-*O*-xylosyl-rutinoside and cyanidin-3-sambubioside, for both WB and E samples were measured by HPLC-MS. The lyophilized samples were dissolved in 80:20 (Water:sample). The solution was filtered and then injected into LC-MS system (Agilent 6538 UHD-QTOF equipped with Agilent 1290 infinity UPLC). Upon extracting the chromatograms based on the reported m/z, calculations were performed by integrating the peak to obtain the area. Anthocyanin standards were purchased from Extrasynthese S.A.S. (Lyon, France).

### *Helicobacter pylori* strains and culture conditions

Two different *cagA*^*+*^, *vacA s1/m1 H. pylori* strains, originally isolated from human patients, were used in our experiments: strain 60190 (kind gift from Dr. G. Perez-Perez, New York University, ATCC #49503) and strain PMSS1 (kind gift from Dr. K. Wilson, Vanderbilt University), which is widely used in murine infection experiments. Bacteria were grown at 37°C under semi-anaerobic conditions on Brucella agar plates, 5% sheep blood (Becton Dickinson) for 3 days. Colonies were harvested into warm Brucella broth supplemented with 10% FBS and were then cultured in a shaking incubator for a further 18 h period prior to use in the experiments.

### High-throughput *Helicobacter pylori* growth assay

High throughput bacterial growth assays were performed in 96-well plates using an OmniLog (Biolog, Hayward, CA) plate reader-incubator. Bacterial growth was visualized with a proprietary redox-sensitive tetrazolium dye [19] (dye D, Biolog). Serial dilutions of berry suspensions or extracts (0.26-4.17% w/v) prepared in IF-10a were added to the plates as indicated together with dye D (0.01%) and PM additive (0.05% BSA, 0.01% NaHCO_3_ and 0.045% glucose w/v final concentrations). Live *H. pylori* were resuspended in IF-10a (Biolog) to a final OD600 = 0.5, which corresponds to 3.4 ×10^8^ bacteria/mL [20], and 20 μL of the bacterial suspension were added to the plates together with berry preparations at appropriate dilutions, for a total volume of 120 μL. For analysis, loaded plates were sealed in a gas-impermeable bag with a CO_2_ Compact sachet (Oxoid, Nepean, ON, Canada) and were incubated at 37 °C in the OmniLog incubator for 30 - 48 h. Absorbance values were recorded at 562 nm every 15 min. In some instances, 10-fold serial dilutions of the *H. pylori* cultures were recovered from the plates and were re-streaked on Brucella agar plates to confirm growth and growth suppression.

### Data analysis for bacterial growth assays

To analyze *H. pylori* growth inhibition by berry compounds, OmniLog data were exported to Excel using the Biolog Data Converter and PM Analysis Software (Microbe) (Biolog, Hayward, CA). Absorption data for each sample and time point were normalized to baseline by subtracting the average of the first four absorption values from each data point. To quantitate bacterial growth over time, peak area under the curve (AUC) was determined using GraphPad version 8.3.1 (San Diego, CA). The minimum inhibitory concentration 90 (MIC_90_) of a berry preparation was defined as the concentration were the AUC values for the growth curves were decreased to ≤10 % of the maximum.

### Human gastric organoid culture and viability assay

Human gastric organoid cultures (gastroids) were established and maintained as previously described [21, 22]. Briefly, human gastric tissue samples were obtained with informed consent and IRB approval from patients undergoing endoscopy and biopsy at the Bozeman Health Deaconess Hospital (protocol DB050718-FC). Alternatively, tissue samples from sleeve gastrectomy surgeries were provided by the National Disease Research Interchange (protocol DB062615-EX). None of the donors were positive for active *H. pylori* infection, as determined by rapid urease CLO test. Gastric glands were prepared by collagenase digestion and then were plated in Matrigel. Following polymerization, Matrigel was overlaid with L-WRN medium which includes Advanced DMEM/F12 (Gibco by Life Technologies, Grand Island, NY) and 50% supernatant from murine L-WRN cells—which secrete Wnt3a, noggin, and R-spondin 3—and supplemented with 10% FBS (Rocky Mountain Bio, Missoula, MT), 1% L-Glutamine, 10μM Y-27632 (Tocris Biosciences, Bristol, UK), 10 μM SB-431542 (Tocris Biosciences, Bristol, UK), and 10mM HEPES buffer. Black raspberry extract prepared in 90% DMSO and 10% HCl was externally administered to the organoids for a 48-hour treatment at 37°C with 5% CO_2_. To determine cell viability, organoids were harvested by trypsinization, and single cell suspensions stained with 7-aminoactinomycin D (7-AAD; ThermoFisher Scientific, Waltham, Massachusetts) were analyzed on an LSR II flow cytometer (Becton Dickinson).

### *H. pylori* infection and berry application in mice

Mouse experiments were performed in 6 – 12 week-old male and female C57BL/6 mice housed in the Animal Resources Center at Montana State University (MSU) following standard protocols. Experimental protocols were approved by MSU’s Institutional Animal Care and Use Committee (protocol #2018-80). Two weeks before infecting the mice with *H. pylori*, animals were placed on a powdered diet (AIN-93M, Dyets. Inc, Bethlehem, PA). For *H. pylori* infection, the PMSS1 strain was prepared as described above. Bacterial numbers were determined by spectrophotometry at 600 nm based on a standard curve generated using the LIVE/DEAD BacLight Bacterial Viability and Counting Kit (Molecular Probes, Eugene, OR), as previously described [20, 23]. Mice were gavaged with a suspension of 10 × 10^8^ *H. pylori* bacteria in 0.1 mL of Brucella broth per mouse twice over 3 days. Two weeks after the first *H. pylori* inoculation, mice were switched to a powdered diet containing 10% w/w of black raspberry powder (BerriHealth, BH-BRB), or a calorie-matched diet containing 10% corn starch. Four weeks later, mice were euthanized with isoflurane followed by cervical dislocation, and stomachs were aseptically harvested for analysis. To quantify bacterial infection rates, gastric tissue homogenates were serially diluted and then plated on TSA plates supplemented with 5% sheep’s blood, 20 μg/mL vancomycin, 30 μg/mL bacitracin, 10 μg/mL nalidixic acid, and 2 μg/mL amphotericin for colony counting. Pathological alterations of the stomach were assessed on H&E-stained paraffin-embedded tissue sections by a blinded anatomic pathologist (F.M.) using the scoring system described by Roth et al. [24].

### Statistical analysis

Data shown are representative of three or more replicate experiments. For OmniLog assays, 3-6 technical replicates were prepared. All data were analyzed using GraphPad version 8.3.1 (San Diego, CA). Data area shown as mean ± SD. The Student’s *t* test or a one- or two-way ANOVA with Tukey’s or Dunnett’s multiple comparisons test was used to determine statistical significance. Differences were considered significant at *P* ≤ 0.05.

## Results

### Powders and organic extracts of black raspberries and blackberries contain variable amounts of anthocyanins

In order to study the potential antibacterial effects of black raspberry (BRB), red raspberry (RRB) and blackberry (BB) compounds on *H. pylori*, we first prepared suspensions and extracts of berry powder, using either fresh-frozen berries or freeze-dried powders from different suppliers and origins as starting materials, as shown in **Table 1**. One additional BRB extract was kindly provided by Dr. S. Hecht, University of Minnesota.

Amongst the multiple bioactive natural compounds, anthocyanins in colored berries of the genus *Rubus* have attracted special attention. Anthocyanins are glycosylated, water-soluble phenolic compounds that are responsible for red, purple and blue coloring of multiple berry species, including RRB, BRB and BB [14]. Anthocyanins are strong antioxidants that have been used successfully in cancer chemoprevention models [25] and that also have been implicated in the antibacterial activities of berry preparations [26, 27]. To determine the concentration of anthocyanins, all samples were analyzed by LC/MS for the presence of keracyanin (cyanidin-3-*O*-rutinoside), kuromanin (cyanidin-3-*O*-glucoside) and cyanidin-3-*O*-xylosyrutinoside (**Fig. 1 A,B**). Because of overlapping peaks, the cyanidin-3-*O-*xylosyrutinoside may include cyanidin-3-*O-*sambubioside, another phenolic berry compound that has a similar composition and MW as the xylosylrutinoside, and that is known to be present in BRB at a low concentration [28].

Total anthocyanin content (TAC) was calculated by adding up the concentrations of all detected anthocyanins (**Table 1** and **Fig. 1C**). Overall, large variations in anthocyanin concentrations were observed for berries from different sources and with different processing techniques. Interestingly, lyophilized but otherwise untreated berry powder from RRB and BB contained significantly higher amounts of anthocyanins than the water/ethanol extracts that we had prepared (**Fig. 1C**). This was likely due to an inefficient recovery of anthocyanins in extracts prepared from fresh-frozen berries that were lyophilized in-house (**Fig. 1D**, *P*<0.001, Student’s *T* test). Anthocyanin recovery was higher if extracts were prepared from commercial berry powders (**Fig. 1D**). Individual data for cyanidin-3-*O*-glucoside, cyanidin-3-*O*-rutinoside, and cyanidin-3-*O*-xylosylrutinoside are presented in **Table 2.** Notably, BRB, RRB and BB powders contained similar levels of cyanidin-3-*O*-sambubioside and cyanidin-3-*O*-glucoside, but cyanidin-3-*O*-rutinoside levels were significantly higher in BRBs than in RRBs and BBs, as previously described (**Fig. 1E**, *P*<0.05, mixed model ANOVA).

**Table 2:**
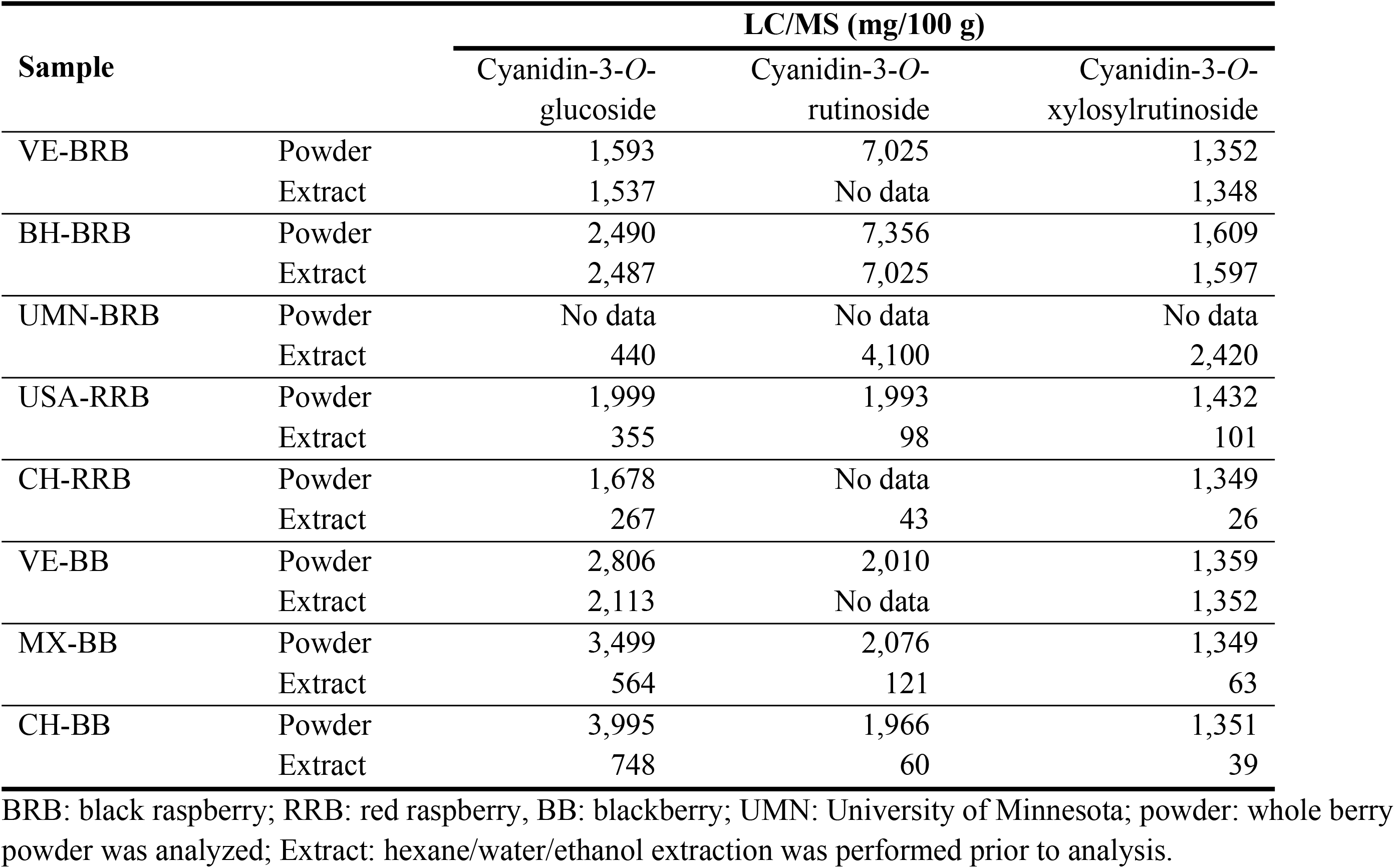
Anthocyanin composition within powdered berries and berry extracts determined by LC/MS.

| Sample |  | LC/MS (mg/100 g) |  |  |
| --- | --- | --- | --- | --- |
|  |  | Cyanidin-3-*O*-glucoside | Cyanidin-3-*O*-rutinoside | Cyanidin-3-*O*-xylosylrutinoside |
| VE-BRB | Powder | 1,593 | 7,025 | 1,352 |
|  | Extract | 1,537 | No data | 1,348 |
| BH-BRB | Powder | 2,490 | 7,356 | 1,609 |
|  | Extract | 2,487 | 7,025 | 1,597 |
| UMN-BRB | Powder | No data | No data | No data |
|  | Extract | 440 | 4,100 | 2,420 |
| USA-RRB | Powder | 1,999 | 1,993 | 1,432 |
|  | Extract | 355 | 98 | 101 |
| CH-RRB | Powder | 1,678 | No data | 1,349 |
|  | Extract | 267 | 43 | 26 |
| VE-BB | Powder | 2,806 | 2,010 | 1,359 |
|  | Extract | 2,113 | No data | 1,352 |
| MX-BB | Powder | 3,499 | 2,076 | 1,349 |
|  | Extract | 564 | 121 | 63 |
| CH-BB | Powder | 3,995 | 1,966 | 1,351 |
|  | Extract | 748 | 60 | 39 |

BRB: black raspberry; RRB: red raspberry, BB: blackberry; UMN: University of Minnesota; powder: whole berry powder was analyzed; Extract: hexane/water/ethanol extraction was performed prior to analysis.

### Development and validation for a high-throughput assay to measure *H. pylori* growth

To determine whether BRB, RRB and BB extracts have antibacterial effects against *H. pylori*, we developed a metabolic bacterial growth assay using the BioLog® system. This system enables kinetic analysis of microbial growth in a 96-well format based on detection of a redox-sensitive dye by the OmniLog® incubator-reader [29]. Since optimal *H. pylori* growth requires microaerophilic conditions, the 96-well plates were sealed into a plastic sleeve with a CO_2_Gen Compact sachet to reduce oxygen levels. As shown in **Fig. 2A** and the **Supplemental video**, addition of *H. pylori* bacteria to the plates at different dilutions resulted in a dose-dependent color response over 48 h. Growth curves had a typical appearance, with an exponential growth phase followed by a plateau phase (**Fig. 2B**). Area under the curve measurements showed significant differences in the growth of *H. pylori* plated at different concentrations, which was confirmed by endpoint measurements at 590 nm in a standard ELISA reader (**Fig. 2C,D**). These results show that *H. pylori* growth can be effectively analyzed in liquid cultures using a high-throughput metabolic growth assay.

**Figure 2.**
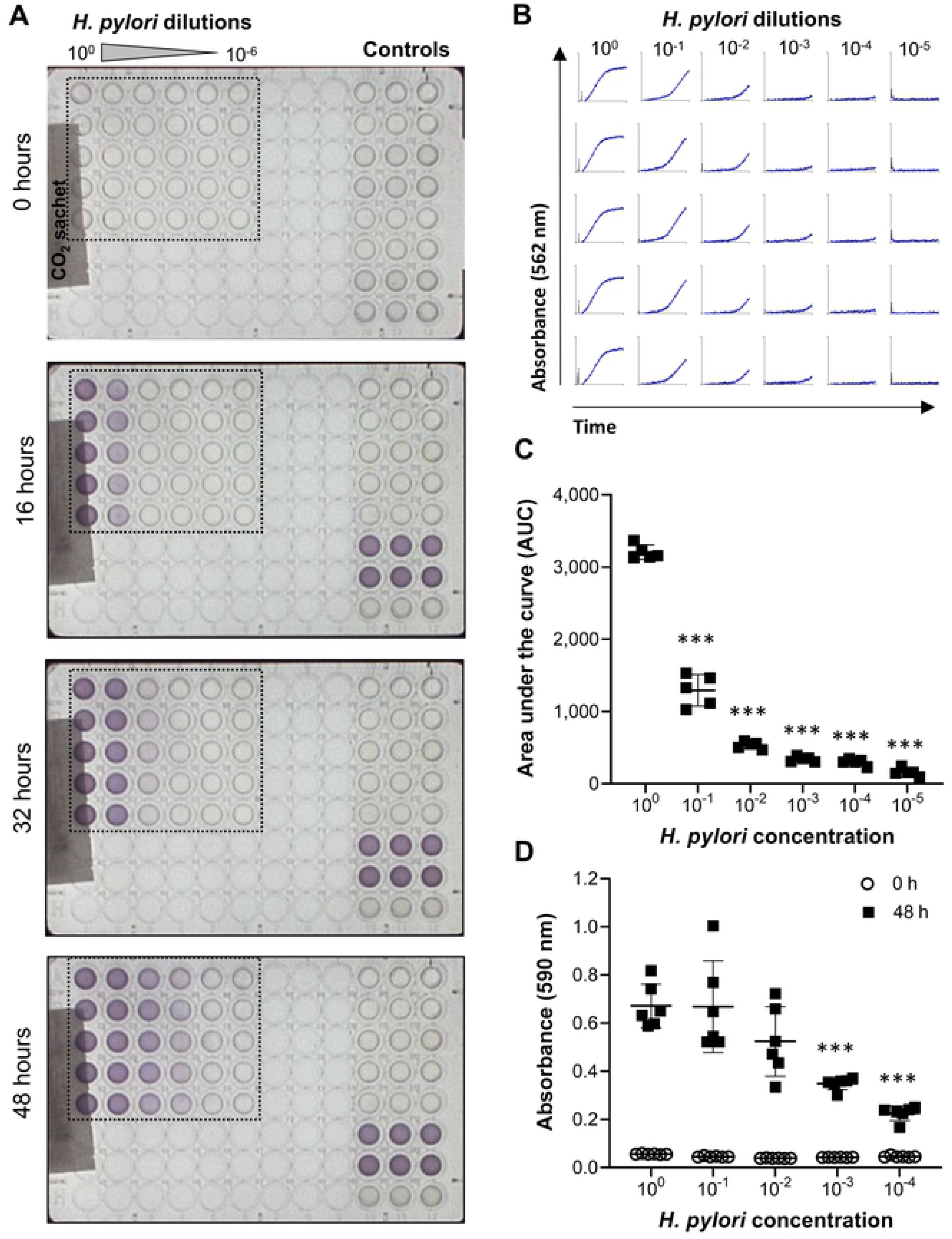
Development of a high-throughput metabolic assay to measure *H. pylori* growth. **(A)** Images of 96-well plates containing various concentrations of *H. pylori* (10^0^ = stock solution used at OD_600_ = 0.5) obtained by the OmniLog incubator/reader at different time points after plating the bacteria. (**B**) Growth curves based on absorbance at 562 nm for the wells outlined in panel A. (**C**) Area under the curve was determined using GraphPad Prism and shows significant differences in *H. pylori* metabolism between cultures with different initial concentrations of bacteria. (**D**) End point absorbance (48 h) of an *H. pylori* culture analyzed in a standard 96-well plate reader at 590 nm. Data are representative of n=4 similar experiments with 5-6 technical replicates each. Individual datapoints, mean ± SD are shown. *** *P*< 0.01 compared to undiluted bacteria, one-way ANOVA with Dunnett’s multiple comparisons.

**Supplemental video.** Time lapse series of plate images of the *H. pylori* culture in Fig. 2, obtained on the OmniLog incubator-reader, shows color development consistent with *H. pylori* metabolism and growth over 48 h.

### The OmniLog® high throughput metabolic assay reveals antibacterial effects of black raspberries on *H. pylori*

We next used the metabolic growth assay to determine whether anthocyanin-rich berry extracts would inhibit *H. pylori* growth. First, we confirmed that the colored berry extracts did not interfere with dye detection in the redox assay. To correct for baseline absorption due to berry compounds, average absorbance over the first four time points measured was subtracted from absorbance measured at each time point during the 48 h experiment. As shown in **Fig. 3A**, BRB powder (4.17%) caused no significant absorption over baseline after 48 h, whereas in the presence of both *H. pylori* PMSS1 and the metabolic dye, significant absorbance was measured (*P*≤0.001). Absorbance was significantly decreased in the presence of BRB powder. Growth curves over 30 h revealed a concentration-dependent inhibition of *H. pylori* growth at BRB powder concentrations between 0.26% and 4.17% (**Fig. 3B**). Area under the curve (AUC) calculations for the *H. pylori* growth curves similarly showed a significant, concentration-dependent decrease of bacterial growth beginning at 0.26% of berry powder (**Fig. 3C**). To validate the metabolic growth data, 48 h liquid cultures from the growth experiments were re-plated on Brucella agar plates and analyzed for formation of *H. pylori* colonies. Consistent with the results from the metabolic assay, complete growth inhibition was seen at 2.08% and 4.17% of BRB powder, demonstrating strong antibacterial activity of the blackberries on *H. pylori*, whereas colony formation was observed in the absence of BRBs or with lower BRB concentrations (**Fig. 3C**). These results demonstrate the ability of BRBs to inhibit *H. pylori* growth *in vitro* and the effectiveness of the OmniLog assay for evaluating *H. pylori* growth and growth suppression by berry compounds.

**Figure 3.**
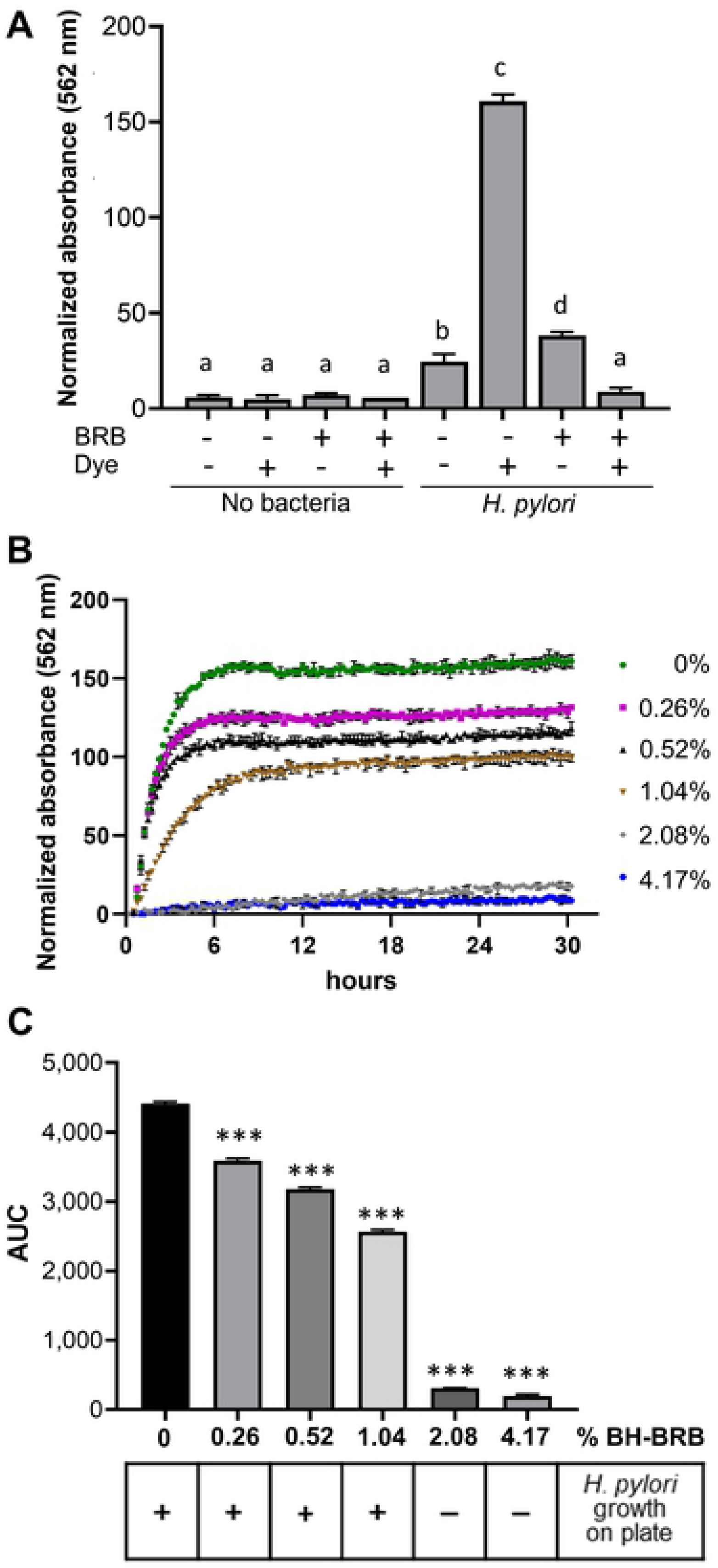
Utilization of the OmniLog microarray to measure antibiotic effects of black raspberries in liquid *H. pylori* PMSS1 cultures. (**A**) End point absorbance values measured by the OmniLog after 48 h of culture with and without *H. pylori*, 8% BH-BRB powder, and/or tetrazolium dye. One representative experiment of three independent experiments with triplicate wells is shown. One-way ANOVA; different letters indicate statistically significant differences at *P≤*0.001. (**B**) Absorbance of *H. pylori* cultures measured over time in the presence of different concentrations of BH-BRB powder for the experiment shown in A. Mean ± SD of triplicate wells, representative of one out of three experiments. (**C**) Top: area-under-the-curve (AUC) values for the experiment shown in A and B. One-way ANOVA; *** indicates statistically significant differences from the untreated control at *P≤*0.001. Bottom panel: matched *H. pylori* growth data on plates, representative of 4 independent experiments.

### Growth inhibition of *H. pylori* cultures by different berry preparations

As shown in Fig. 2 and Tables 1 and 2, composition of berry preparations was highly variable, depending on berry species, source and processing method. Therefore, we next compared the different BRB, RRB and BB powders (whole berries) and extracts described above (Tables 1 and 2) for their ability to inhibit the growth of two *H. pylori* strains, 60190 and PMSS1 in the OmniLog microarray assay. All berry preparations tested had significant antibacterial activity (**Fig. 4 and 5**), with complete inhibition of *H. pylori* growth generally achieved at a concentration of about 4%. The different berry preparations showed a great variation in their ability to suppress *H. pylori* growth. The UMN BRB extract had the strongest antibacterial activity of all the preparations tested, whereas powdered berries generally were less effective. Both *H. pylori* strains had similar responses to the different berry extracts.

**Figure 4.**
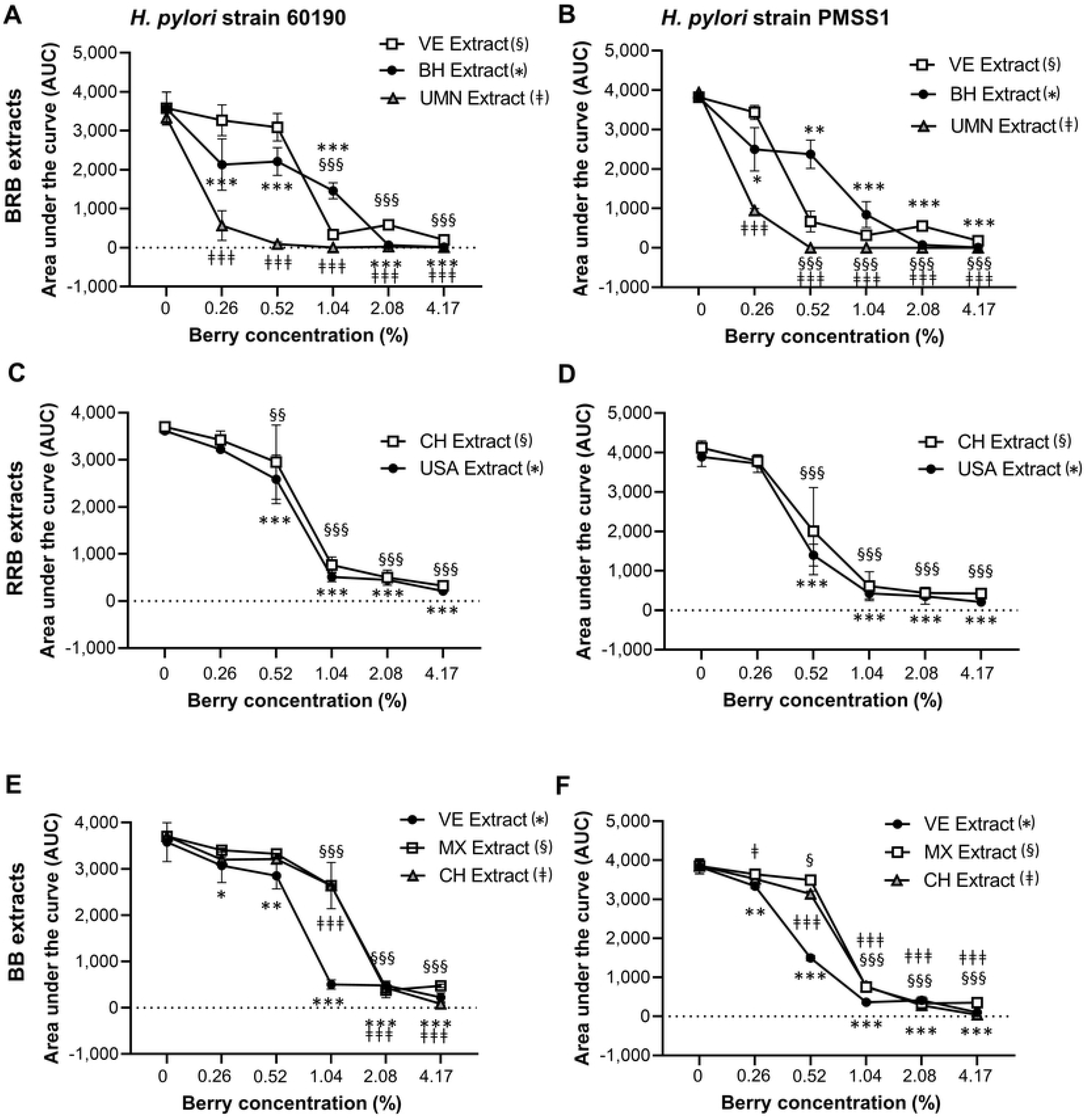
Growth inhibition of *H. pylori* by extracts from BRB, RRB and BB. Growth of *H. pylori* in the presence of different concentrations of the BRB, RRB and BB extracts described in Table 1 and 2. Area under the curve (AUC) for metabolic culture activity was measured over 30 h for strain 60190 with (**A**) BRB extracts, (**C**) RRB extracts and (**E**) BB extracts and for strain PMSS1 with (**B**) BRB extracts, (**D**) RRB extracts and (**F**) BB extracts using the OmniLog assay. Pooled data from n=2-4 independent experiment with 3 technical replicates are shown. Data were analyzed by 2-factorial ANOVA. Statistically significant differences from the untreated control at *P≤*0.05/*P≤*0.01/*P≤*0.001, respectively, are indicated by */**/***, §/§§/§§§ and ǂ/ǂǂ/ǂǂǂ as defined in the panel labels.

**Figure 5.**
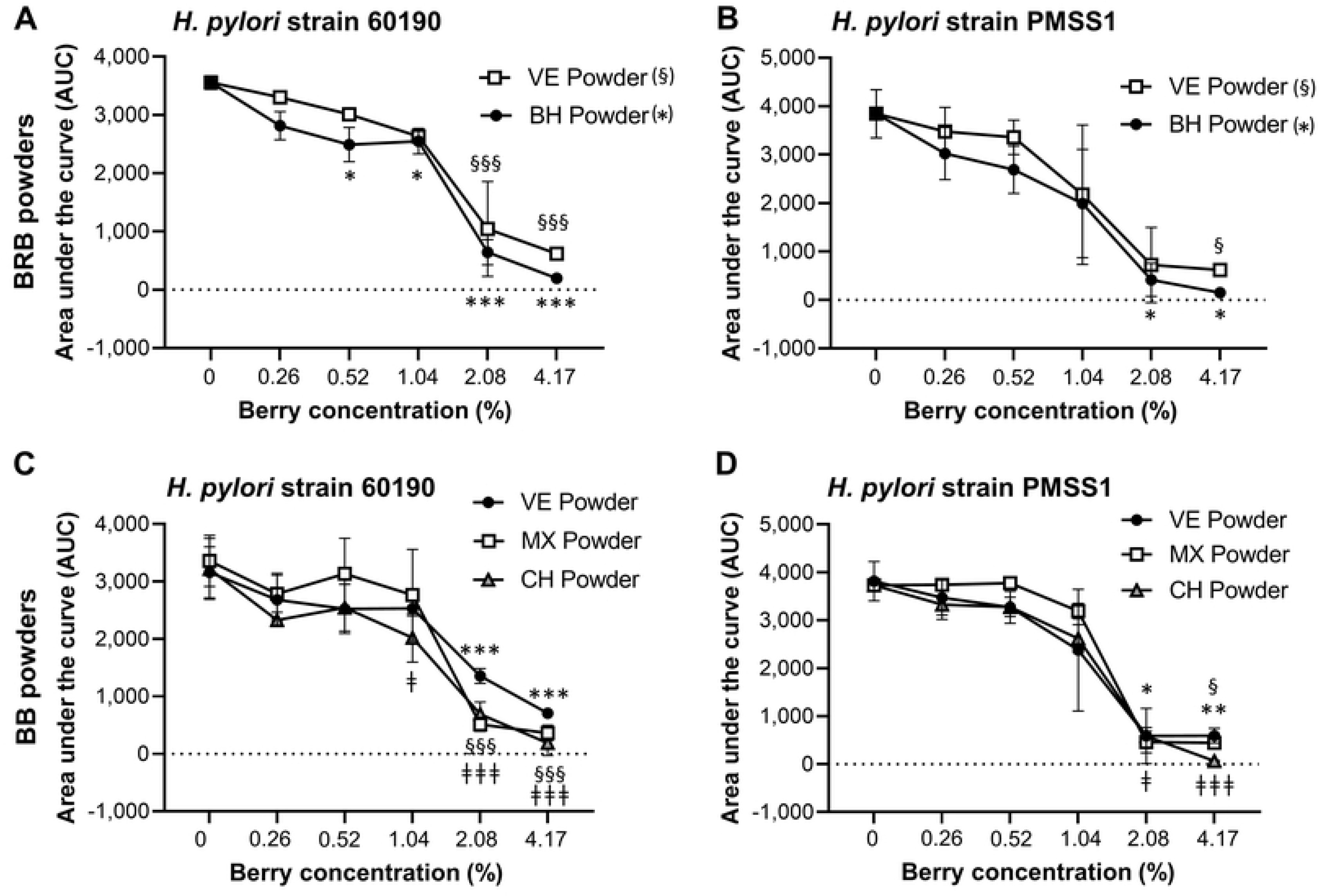
Growth inhibition of *H. pylori* by lyophilized BRB and BB powders. (**A**) Growth of *H. pylori* strains 60190 and PMSS1 in the presence of different concentrations of the BRB and BB powders described in Table 1 and 2. Area under the curve (AUC) for metabolic culture activity was measured over 30 h for strain 60190 with (**A**) BRB extracts and (**C**) BB extracts and for strain PMSS1 with (**B**) BRB extracts and (**D**) BB extracts using the OmniLog assay. Pooled data from n=2-4 independent experiment with 3 technical replicates are shown. Data were analyzed by 2-factorial ANOVA. Statistically significant differences from the untreated control at *P≤*0.05/*P≤*0.01/*P≤*0.001, respectively, are indicated by */**/***, §/§§/§§§ and ǂ/ǂǂ/ǂǂǂ as defined in the panel labels.

To better understand the large variability in the antibacterial effects of the different berry preparations, we performed multifactorial analysis of variance (ANOVA) of the minimum inhibitory concentrations (MICs) required to suppress *H. pylori* growth (**Fig 6A**). Extracts exhibited a stronger antibacterial response, as evidenced by significantly lower MICs (*P≤*0.001). For BRB and BB, the type of berry preparation used (powdered lyophilized powder vs. extract) was responsible for 23% of the variation in MIC. The type of berry (BRB, RRB or BB) and the strain of bacteria also significantly impacted MIC, with BRB associated with lower MICs than BB (*P=*0.019), and strain PMSS1 exhibiting a slightly higher sensitivity to the antibacterial effects of berries than 60190 (*P*=0.039).

**Figure 6.**
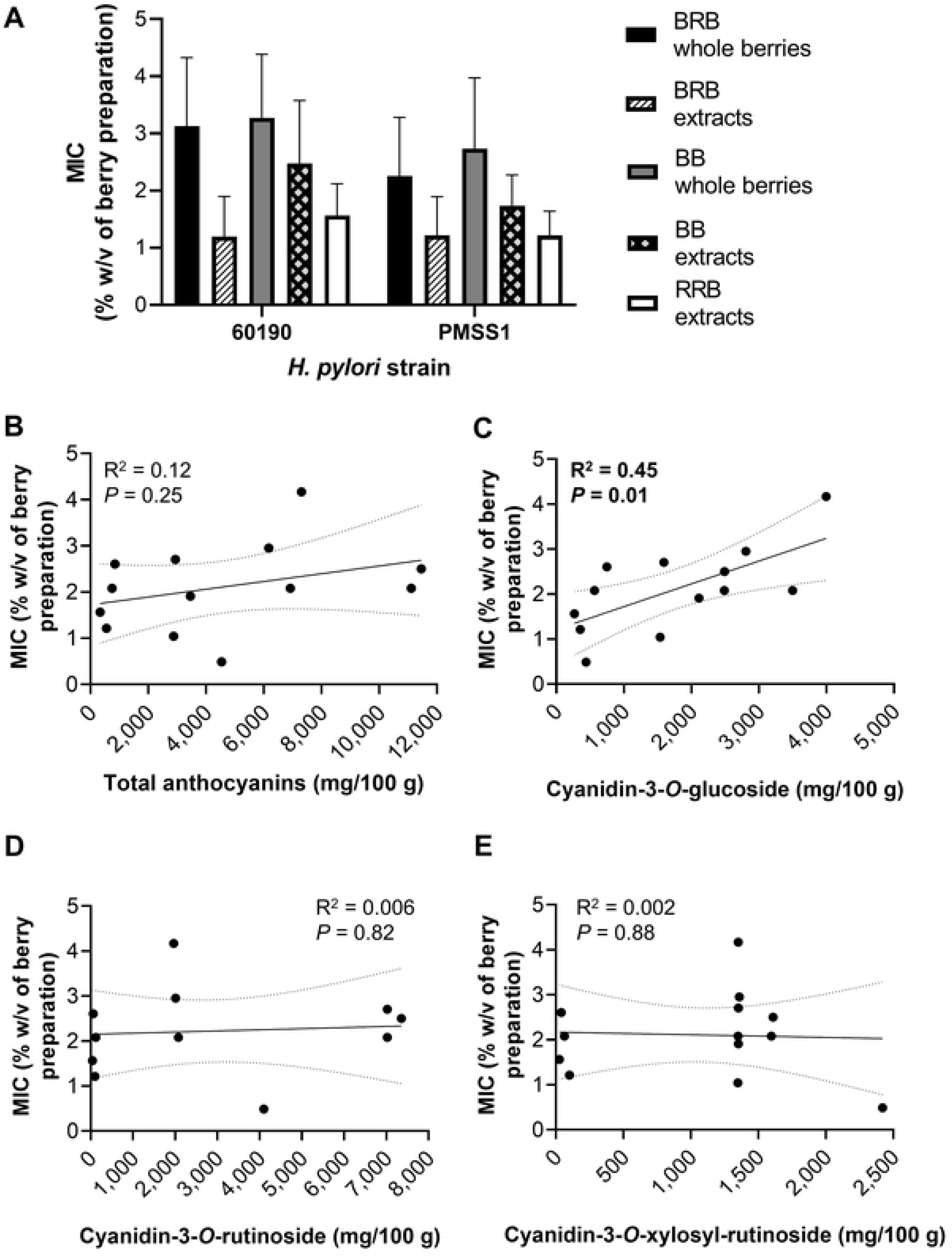
Minimum inhibitory concentrations (MICs) of berry preparations for antibacterial activities against *H. pylori*. (**A**) MIC means ± SD for different berry types and preparations against *H. pylori* strains 60190 and PMSS1. (**B-E**) Correlation between mean MIC observed with both *H. pylori* strains and concentrations of (**B**) total anthocyanins, (**C**) cyanidin-3-*O*-glucoside, (**D**) cyanidin-3-*O*-rutinoside and (**E**) cyanidin-3-*O*-xylosylrutinoside for all berry preparations analyzed. R^2^ = Pearson’s correlation coefficient.

Since anthocyanins are considered the major active ingredients of black and red berries, we hypothesized that anthocyanins would also be responsible for the antibacterial activities observed in our experiments. Therefore, we anticipated that correlation analysis (Pearson’s correlation coefficient) would show a significant inverse relationship between the anthocyanin concentration and the MIC of the berry preparations. However, surprisingly, no significant relationship between the MIC and the total anthocyanin contents or the concentration of cyanidin-3-*O*-rutinoside or cyanidin-3-*O*-xylosylrutinoside was detected (**Fig. 6B, D, E**). In contrast, there was a significant positive correlation between cyanidin-3-*O*-glucoside concentrations and the MICs (**Fig. 6C**, *P*=0.01, R^2^=0.45), indicating that high concentrations of cyanidin-3-*O*-glucoside prevented antibacterial activities. These finding suggest that berry components other than the three anthocyanins analyzed contribute to antibacterial activities of BRB, BB and RRB against *H. pylori*.

### BRB extract demonstrates a high biocompatibility in a human gastric organoid model

Having shown significant antibacterial effects of multiple different berry preparations against *H. pylori in vitro*, we next sought to confirm that the berries were not toxic to the gastric epithelium, where *H. pylori* bacteria generally reside. We used human gastroids (gastric organoids) as a model of primary human gastric epithelial cells, since they closely represent the architecture and cellular complexity of the human gastric mucosa [30, 31]. Organoids are 3-dimensional permanent cultures of primary cells maintained in an extracellular matrix gel in the presence of specific growth factors. Five organoid lines derived from non-*H. pylori-*infected human gastric biopsies or surgical material were cultured in the presence of UMN-BB extract, which had the greatest antibacterial activity of all berry preparations tested (**Fig. 4**). As shown in **Fig. 7**, the organoids tolerated the BRB extract over a wide range of concentrations up to 5 mg/mL (0.5%) without any significant impact on organoid viability, as determined by flow cytometric analysis of 7-AAD staining and phase contrast microscopy. Importantly, the UMN-BRB extract achieved significant inhibition of *H. pylori* growth at a concentration of 0.26% with both *H. pylori* strains tested.

**Figure 7.**
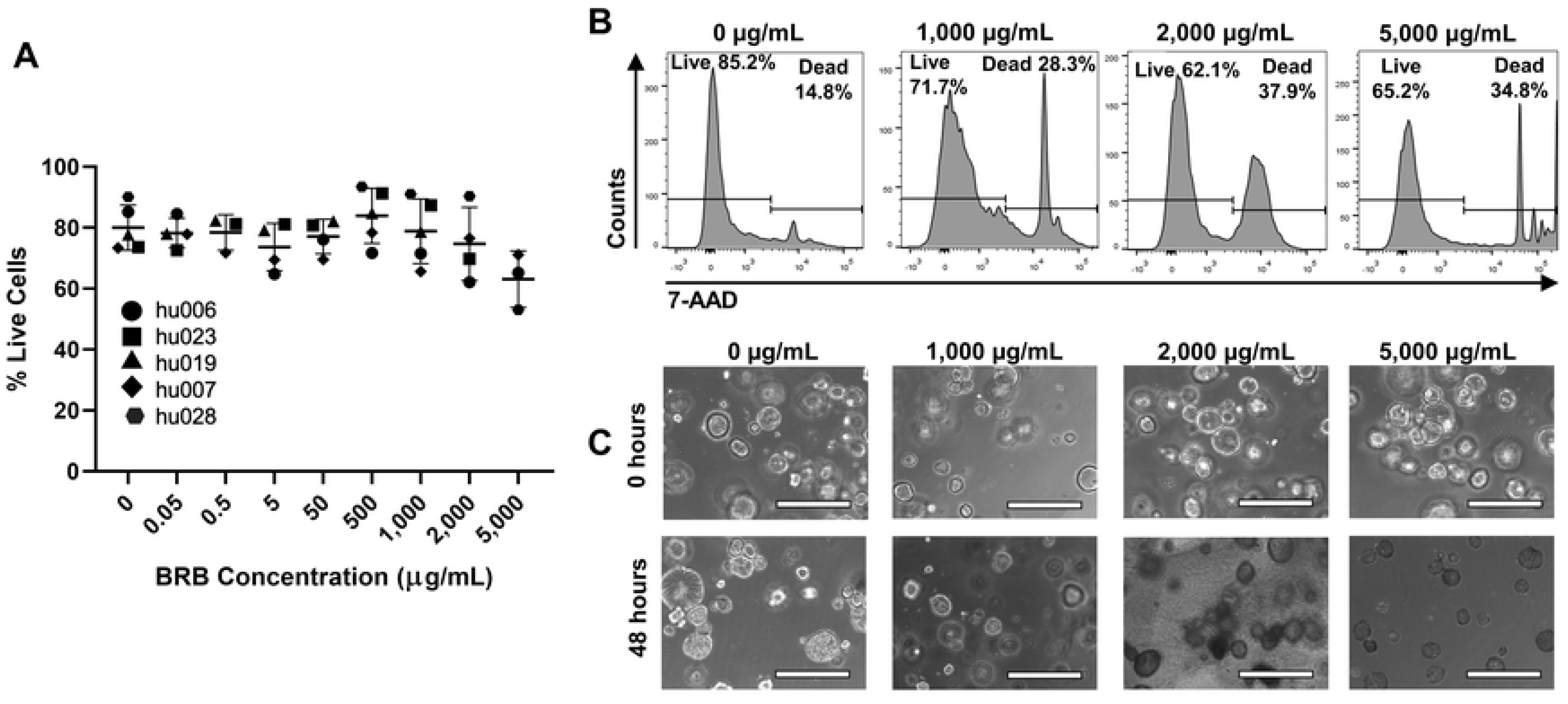
Toxicity analysis of BRB extract in primary human gastric epithelial cell cultures. (**A**) Gastric organoid viability following 48 h of culture in the presence of UMN-BRB extract at different concentrations. Following BRB exposure, cells were harvested by trypsinization and analyzed for 7-AAD dye exclusion by flow cytometry. Pooled data from n=3-5 independent experiments; individual data point, mean ± SD. Differences between treated organoids and untreated controls were analyzed by ANOVA. (**B**) Representative FACS histograms for the data in A showing 7-AAD staining of gastric organoid cells. (**C**) Representative images of organoid cultures treated with UMN-BRB extract after 0 h and 48 h. Bar = 400 μm.

### Dietary BRB powder does not prevent *H. pylori* infection and gastric pathology in C57BL/6 mice

To evaluate whether the antibacterial effects of berries against *H. pylori* observed *in vitro* can protect from *H. pylori* infection *in vivo*, we performed an *H. pylori* infection experiment in C57BL/6 mice. Mice were adapted to a powdered diet for 2 weeks and then were infected with *H. pylori* PMSS1 by oral gavage. Two weeks after the infection, mice were switched to a diet containing 10% of BH-BRB powder, chosen because of its high anthocyanin content, or 10% starch. This application protocol has been successfully used in our previous chemoprevention studies for gastrointestinal tract cancers [32]. Mice were sacrificed four weeks after initiation of the berry diet (**Fig. 8A**), and stomachs were analyzed for the presence of *H. pylori* and scored for pathological alterations. As shown in **Fig. 8B**, all mice were colonized with *H. pylori* at the end of the experiment, with no significant difference between the group fed the berry diet compared to the group fed the control diet. Furthermore, *H. pylori* infection led to significant gastric inflammation, but this was not significantly altered by the absence or presence of dietary berries.

**Figure 8.**
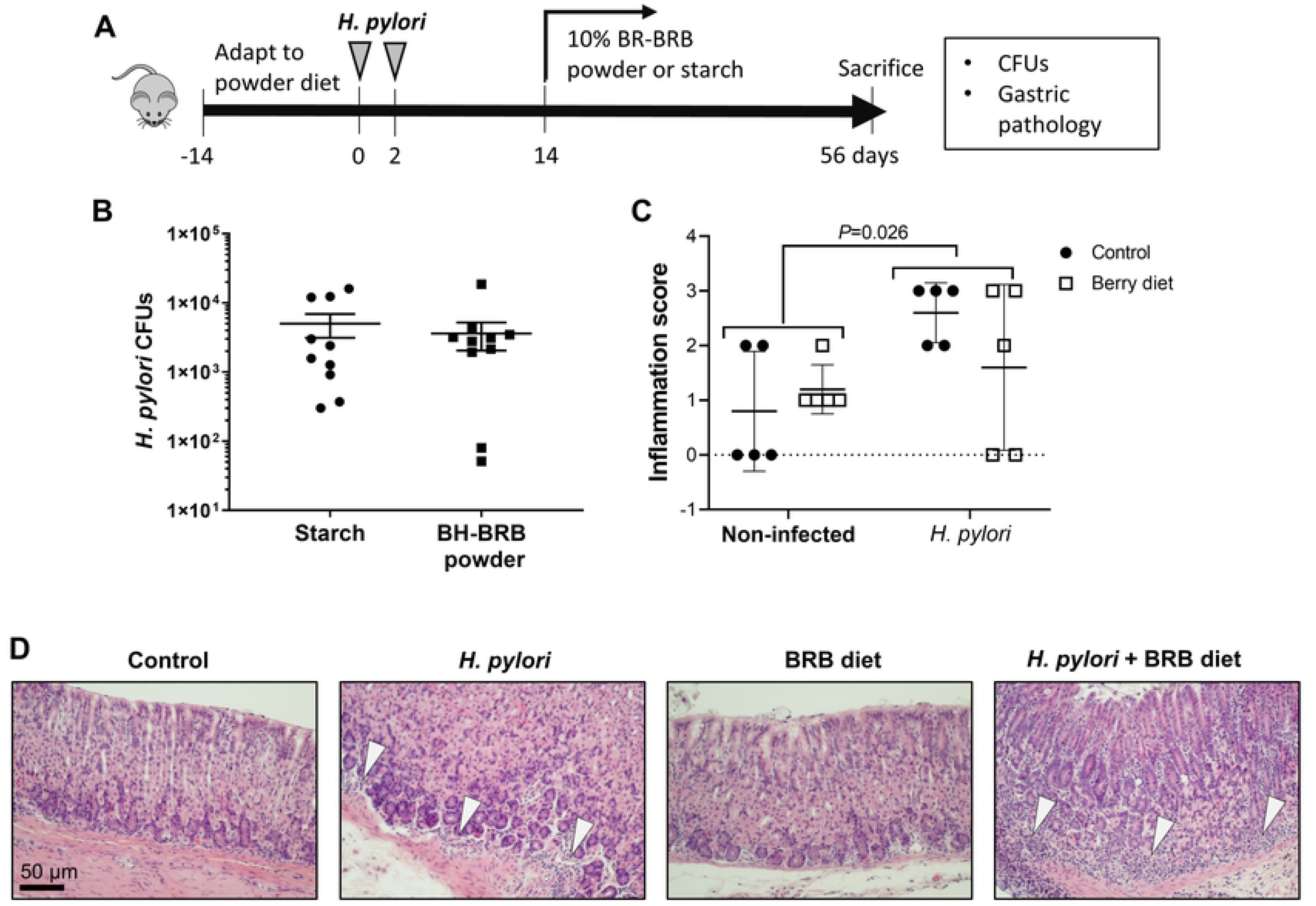
Dietary application of 10% BRB powder in C57BL/6 mice does not decrease gastric *H. pylori* colonization or gastric pathology. **(A)** Experimental design. **(B)** Gastric *H. pylori* colonization levels were determined by colony counts. Pooled data from two independent experiments with n=5 six-to twelve-week-old male and female C57BL/6 mice each. Individual data points, mean ± SD. Differences between the groups were analyzed by Student’s *t* test. (**C**) Analysis of gastric pathology following *H. pylori* infection and dietary berry application. Individual data points, mean ± SD, n=5 per group. Data were analyzed by two-way ANOVA with Tukey’s multiple comparisons test. (**D**) Representative images of gastric tissue collected at the end of the experiment shows inflammatory infiltrates in the presence of *H. pylori* infection. Arrowheads point to inflammatory infiltrates. Bar = 50 μm.

## Discussion

In this study, we demonstrated that black and red raspberries and blackberries have significant antimicrobial activity against *H. pylori* in a high throughput bacterial growth assay. Interestingly, our analyses showed that both the chemical composition and antimicrobial activity were highly variable depending on berry type, origin and processing method. In addition, we showed that long-term dietary application of a high concentration of berry powder was insufficient for eliminating *H. pylori* infection in mice.

One major advancement of our study was the development of a high throughput metabolic microarray assay compatible with the growth requirements of the microaerophilic bacterium *H. pylori*, which enabled us to test a large number of different berry preparations at a wide range of concentrations. The efficacy of antimicrobial treatments for *H. pylori* is typically still analyzed using the agar dilution or E strip diffusion methods [33–36], which are work intensive, expensive, and difficult to scale up. In a previous study, Lee et al. utilized proprietary Biolog Phenotype Microarray plates to evaluate the ability of *H. pylori* to metabolize different carbon sources [37]. Here, we used Biolog’s tetrazolium dye and media together with various berry dilutions on standard 96-well plates to dynamically analyze *H. pylori* growth inhibition. By sealing liquid *H. pylori* cultures in transparent, gas-impermeable plastic sleeves with small CO_2_ sachets, microaerophilic growth conditions were maintained. Importantly, microarray culture results matched growth profiles on standard agar plates, as we demonstrated by re-culturing *H. pylori* samples from the microarray plates. The presence of colored berry preparations did not interfere with dye detection, suggesting that our analysis method is suitable for use with other colored natural products as well as a wide range of other chemical compounds.

Antimicrobial properties of *Rubus* berries have been described in multiple previous studies [15, 38–41]. Our analysis of powders and extracts from BRB, RRB and BB from various suppliers and geographical regions revealed that all berry preparations had significant antimicrobial activity against *H. pylori in vitro*, regardless of the *H. pylori* strain used. However, MIC_90_ for the various berry preparations differed significantly, corroborating with data from previous studies. Concentrations of active ingredients in *Rubus* berries are known to vary based on geographical location, environmental conditions and berry type [40, 42]. In addition, post-harvest processing and storage may affect active ingredients [43, 44]. Moreover, different bacterial species vary in their susceptibility to the antibacterial effects of berries [45]. Overall, these previous studies and our data indicate that each berry product needs to be carefully tested for specific biological activity and applications prior to use as a neutraceutical. The top-performing preparation in our study was a black raspberry extract from the University of Minnesota (UMB-BRB-E), which achieved complete *H. pylori* growth inhibition at (>90%) at 0.5 % (5 mg/mL). This inhibitory concentration is similar to or lower than those of *Rubus* extracts described in other studies [16, 38, 46], but several log folds higher than standard antibiotics such as amoxicillin and clarithromycin [47].

Surprisingly, we did not observe any significant correlation between the concentration of any anthocyanin analyzed in our preparations and increased antibacterial activity, indicating that antimicrobial activity was largely independent of anthocyanins. Anthocyanins exhibit antimicrobial activity against gram-negative bacteria by causing damage to the cell walls, membranes, and intercellular matrix [48]. Importantly, anthocyanins have been linked to the antimicrobial effects of berry preparations in previous studies [49–51] and are responsible for the major chemopreventive effects of blackberries and black and red raspberries [25, 28, 52–54]. However, our correlation analysis suggests that the antibacterial effects against *H. pylori* were independent of anthocyanins and thus must be caused by other active compounds. Indeed, ellagic acid, another polyphenol present in red and black berries, is known to exert antibacterial activity against *H. pylori* as well as other bacteria [17, 55, 56]. In addition, Lengsfeld et al. demonstrated that berry-derived polysaccharides can combat *H. pylori* infection *in vivo* by preventing bacterial binding to the gastric mucosa [57]. Additional studies have shown antibacterial effects for berry-derived sanguiin H-6 [41] and rubososide [58]. Further experiments are needed to identify the raspberry and blackberry berry compounds that mediate antimicrobial activity against *H. pylori*.

Although our experiments demonstrated good biocompatibility and significant antimicrobial activity against *H. pylori in vitro*, this did not translate into antibiotic activity in an *in vivo H. pylori* infection model. In previous studies, we have successfully used dietary application of 5% lyophilized powdered BRB to prevent esophageal, oral and colon cancer in rats and colonic polyps in mice [32, 52, 59]. Notably, the present short-term *H. pylori* infection study was not designed to analyze chemopreventive effects of BRB on *H. pylori*-induced gastric cancer, and no significant cancerous or pre-cancerous lesions were observed in any of the animals. While we were unable to demonstrate antimicrobial activity of BRB in mice with our experimental setup, such effects have previously been demonstrated. Thus, Park et al. [17] recently showed that extracts prepared from dried, unripened Korean raspberry (*Rubus crataegifolius)* decreased in *H pylori* colonization by about 4 log-fold in a murine model of infection with *H. pylori* strain SS1. The berry preparation used in the study by Park et al. was highly potent, with an *in vitro* MIC_90_ of 150 μg/ml [17], whereas the MIC90 for the BH-BRB powder that the animals in our study received was 2%, equivalent to 20 mg/mL. Moreover, administering just a single high dose by gavage may have resulted in higher concentrations of active berry ingredients at the gastric epithelial surface, where *H. pylori* resides, than constant dietary ingestion of a lower amount (10%) of berry powder.

In summary, we have established a high throughput metabolic growth assay to analyze antimicrobial effects of berry preparations against *H. pylori.* Both freeze-dried powders and ethanol extracts from BRBs, RRBs and BBs obtained from various sources significantly suppressed growth of multiple strains of *H. pylori in vitro.* Toxicity studies with human gastric organoids demonstrated good biocompatibility over a wide range of concentrations, including the MIC_90_ determined in the growth assay. However, dietary application of BRB powder had no significant effect on *H. pylori* colonization in a mouse model of *H. pylori* infection. Together, our findings confirm the potential of berry products as antimicrobial agents but highlight the importance of optimizing protocols for each berry type, preparation and application.

## Acknowledgments

This study was supported by grant funding from the Oregon Raspberry and Blackberry Commission (to GS and CG; https://oregon-berries.com/), Montana INBRE (NIH award P20GM103474, to DB, AS and KL; https://inbre.montana.edu/) the Montana Agricultural Experiment Station (project #1015768/MONB00450, to DB; https://agresearch.montana.edu/maes.html) and an equipment grant from the M. J. Murdock Charitable Trust (#2016028, to DB; https://murdocktrust.org/grant-opportunities/). The funders had no role in study design, data collection and analysis, decision to publish, or preparation of the manuscript. The collaborative support of Bozeman Health Deaconess Hospital for collecting human tissue samples is greatly appreciated. We also would like to thank Dr. Stephen Hecht, University of Minnesota, for providing BRB extract and Dr. Charlotte Quist, Montana State University, and Dr. Maria Blanca Piazuelo, Vanderbilt University, for assistance with histopathological scoring of murine stomach tissues.

## Author contributions

Study design: CG, GS, and DB; acquisition of data: CG, KL, AM, MMR, FM, TAS, ABG, GB and DB; analysis and interpretation of data: CG, KL, AM, MMR, GB, GS and DB; drafting the article: CG, KL, GB, GS and DB. All authors were involved in revising the manuscript critically for important intellectual content, have approved the final version of the manuscript and have agreed to be accountable for all aspects of the work in ensuring that questions related to the accuracy or integrity of any part of the work are appropriately investigated and resolved.

## Ethics Statement

### Human Subject Research

Human tissue sample collection was approved by the Institutional Review Board of Montana State University, protocols DB050718-FC and DB062615-EX. Written consent was obtained from study participants unless collected materials were de-identified surgical discard materials covered under exempt protocol DB062615-EX.

### Animal Research

Experimental protocols were approved by MSU’s Institutional Animal Care and Use Committee (IACUC), protocol #2018-80. Mice were euthanized following AVMA guidelines using isoflurane inhalation followed by cervical dislocation.

